# An aptamer targeting TfR1 enhances ASO delivery to muscle tissue

**DOI:** 10.64898/2026.07.13.737509

**Authors:** Matthew Warner, Linsley Kelly, Reetu Thakur, Meenakshi Ravichandran, Deepti Tomar, Nidhi Nidhi, Vandana Tamraparni, Elaiyarasu Govindaraj, Priyanka Samji, Mahati Krishna, Abhay Suhas Kulkarni, Matthew Levy

## Abstract

Oligonucleotide based therapeutics continue to rise as a significant class of medicines for the treatment of human disease. However, achieving delivery to non-hepatic tissues remains a challenge in the field. While substantial advances have been realized, largely through the use of antibody or protein based targeting agents to tissues including muscle and the CNS, these large protein agents present complications in synthesis and carry the potential for immune responses. In an effort to identify a simpler, smaller and robust means of delivery, we have generated and evaluated aptamers targeting the human transferrin receptor (hTfR) for the delivery of both ASO and siRNA cargoes to muscle. Using a fully backbone modified anti-TfR aptamer, 36 nt in length, that binds hTfR and does not compete for binding with the natural ligand, transferrin, we evaluated the ability to deliver ASOs to skeletal muscle following systemic delivery. Using optimized linker chemistry, aptamer-ASO conjugates led to >50% target gene knockdown in muscle tissue for up to 42 days following a single dose at 3 mg/kg ASO (∼11 mg/kg total drug) in mice. Taken as a whole, these results offer significant promise for the use of aptamers in the development of future therapeutics.

## Introduction

Targeting the transferrin receptor (TfR1) has now become an established and validated means to drive muscle uptake of oligonucleotide therapeutics including both antisense oligonucleotides (ASOs) and small interfering RNAs (siRNAs). Initially demonstrated using Fab antibody fragments for the delivery of siRNA (1), the approach has been extended to a variety of other delivery platforms and cargoes with the most advanced approaches using full length antibodies showing efficacy in the clinic (2). Similar advances using both protein based (Fab, modified Fcs, etc.) and peptide based approaches to target TfR1 for the delivery of ASOs have also been demonstrated (3–5). However, several challenges remain when using antibody based, protein and even peptide based delivery systems. For antibodies and other proteins, compound development requires a multi-step process including complex biological production and the site-specific conjugation of a polyanion which can complicate production; drug-to-antibody ratios must be carefully controlled (6). Even for peptides, where site-specific handles can be incorporated, their conjugation to oligonucleotides remains a multi-step procedure requiring a post-synthetic conjugation and several purification steps.

Aptamers represent an alternative class of ligand for the targeted delivery of oligonucleotide based therapeutics. As oligonucleotides themselves, these ligands can be appended to oligonucleotide cargoes in “cis”. Targeted ASOs can be generated using standard solid phase chemistry and a single synthesis run; a typical aptamer-ASO conjugate is expected to be <60 nt in length (∼40 for the aptamer and 16 - 20 for the ASO). The conjugate can then be easily cleaved, deprotected and purified using standard approaches. Moreover, because the resulting conjugates are composed solely of nucleic acids using chemistries similar to clinically approved siRNA based drugs, immune-related adverse events are minimal. Indeed, oligonucleotide based drugs such as siRNA have demonstrated very high safety and low ADA rates (7).

To date, the use of aptamers for targeted delivery has had a bit of a sorted past with many compounds failing to robustly function or bind their targets in simple, well-controlled in vitro assays (8). Indeed, when we previously evaluated a selection of 15 well described cell surface targeting aptamers, only 3 of the compounds proved capable of engaging their intended targets (9).

Here, we have developed and explored the use of a fully backbone-modified aptamer which targets the human transferrin receptor and does not compete with the natural ligand, transferrin (Tf), for binding. Using this ligand we generated aptamer-ASO conjugates, 56 nucleotides in length, and evaluated their potential to target and enhance delivery to muscle tissue. This approach greatly simplifies the development of targeted therapeutics and demonstrates that robust aptamers are viable agents for extrahepatic targeted drugs.

## Results

### Evaluation of anti-hTfR aptamers

We performed validation studies to compare the binding properties of some of the published anti-TfR aptamers as well as a fully modified anti-TfR aptamer developed in-house to assess their potential for drug development. Candidate compounds included four different 2’F modified RNA aptamers, c2.min, E3, Waz and TR14ST1-3, and the fully backbone modified aptamer, CR.5 (10–14). Binding was assessed by flow cytometry using fluorescently labeled aptamers and Jurkat cells, a human immortalized T cell line that expresses high levels of transferrin receptor. Assays were conducted in full media, and compounds were incubated with cells for 1 hr at 37°C allowing time for the aptamers to both bind the cells and be internalized. The resultant signal is a combination of these two events. Consistent with literature reports, both c2.min and E3 both showed robust cell binding as assessed by flow cytometry in the absence of the natural ligand, Tf (13, 15). However, in the presence of 12.5 µM Tf, the aptamers lost almost all binding activity (**Fig. 1A**). In contrast, both Waz and CR5 demonstrated robust Tf independent binding. Surprisingly, TR14 ST1-3, a 2’F modified aptamer selected against human TfR, which is reported to compete with Tf for binding to TfR1 and deliver oligonucleotide cargoes to the CNS of wildtype mice following systemic, iv, administration, failed to show any activity in both the presence or absence of Tf (TR14 ST1-3; **Fig. 1A**) (Yoon et al. 2019).

**Figure 1:**
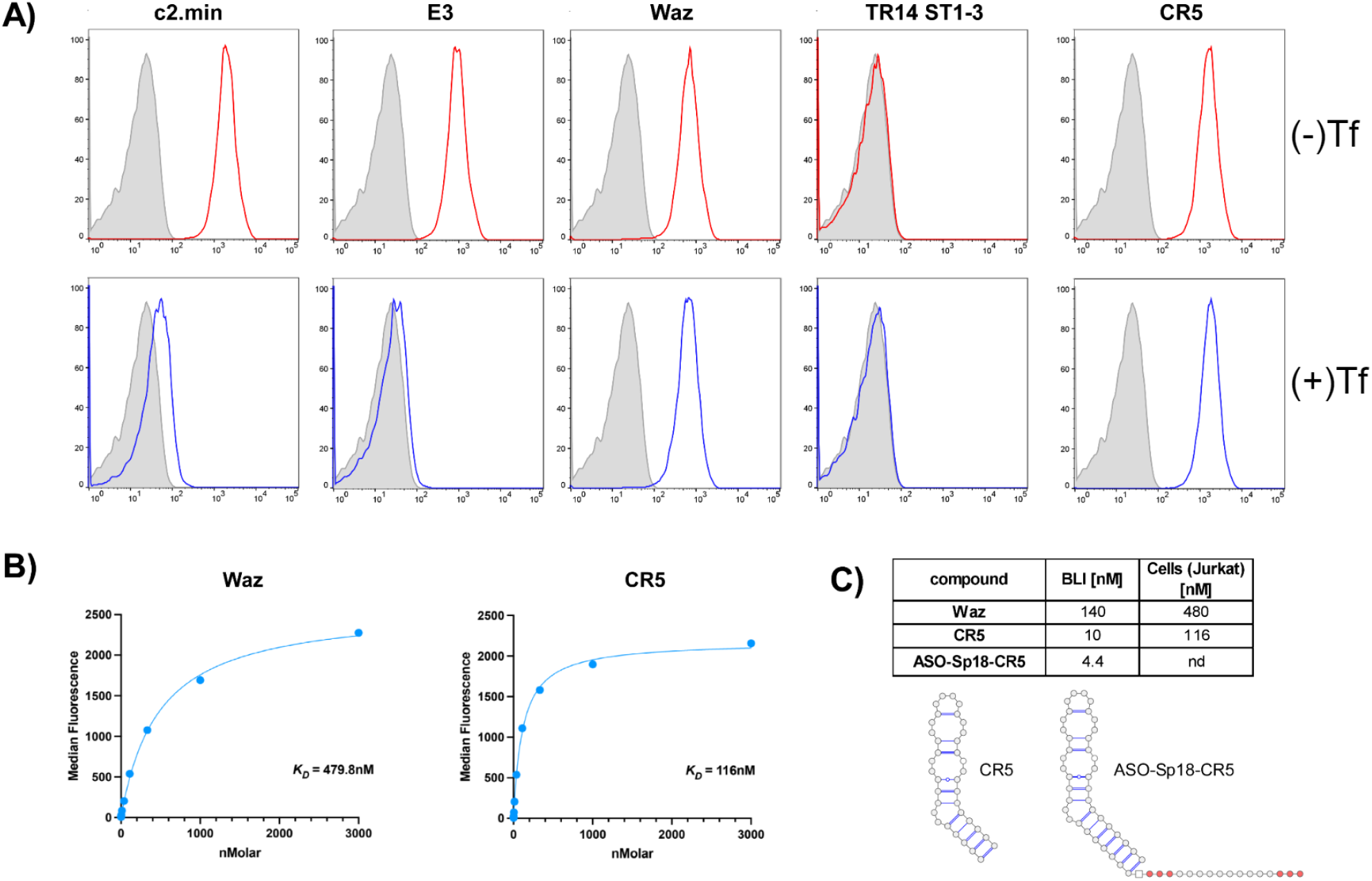
Evaluation of TfR Aptamers. **(A)** TfR aptamers were evaluated by flow cytometry on human Jurkat cells in the presence and absence of 12.5 µM human holo-transferrin. Assays were performed in full media with 200 nM aptamer. **(B)** Dose response analysis of Tf-independent TfR aptamer Waz and CR5. **(C)** Binding affinities as determined by BLI for Tf-independent TfR aptamer Waz, CR5, and the aptamer-ASO conjugate ASO-SP18-CR5.

For ligands like c2.min and E3, competition with the natural ligand, Tf, would likely create toxicity complications for drug development. We therefore chose to focus additional studies on Waz and CR5. As shown in **Fig. 1B**, a dose response analysis on Jurkat cells revealed CR5 binds and is internalized by cells with an apparent IC50 ∼4 fold better than Waz. When we assessed binding affinity using BLI, the difference in affinity was even greater with CR5 demonstrating a K_d_ of 10 nM, compared with 140 nM for Waz (**Fig. 1C**). The better binding characteristics combined with the fully modified backbone composition of CR5 make it the better choice for developing a new targeted oligonucleotide drug. We therefore generated an aptamer-ASO conjugate (ASO-Sp18-CR5) in which the aptamer portion and a 3/10/3 LNA-gapmer targeting DMPK (16) were separated by a hexadecyl ethyleneglycol spacer (Sp18), thus ensuring the aptamer could still function with an appended cargo. The ASO was attached to the 5’ end of the aptamer to avoid complications with desulfurization during synthesis. As shown in **Fig. 1C**, when compared to the parent aptamer, CR5, the ASO conjugate showed similar binding characteristics as assessed by BLI (K_d_ ∼ 4.4 nM).

### In vivo evaluation of anti-TfR aptamer conjugates

TfR targeted aptamer-ASO conjugates were evaluated for activity following iv administration in human transferrin receptor knock-in mice. In short, mice were treated with the aptamer conjugate (ASO-Sp18-CR5) or ASO only at 1, 3, and 10 mg/kg (mass based on ASO) on days 1, 5, and 9. Animals were subsequently sacrificed on day 16 (7 days post last dose) and DMPK knockdown evaluated in the gastrocnemius anterior (GA) muscle and heart. As shown in **Fig 2A**. and **Fig. 2B**, all doses of the aptamer-ASO conjugate led to an increase in target gene knockdown when compared to animals treated with the non-targeted ASO only. Consistent with other studies targeting TfR, a more robust effect was observed in GA muscle versus heart (4, 16).

**Figure 2:**
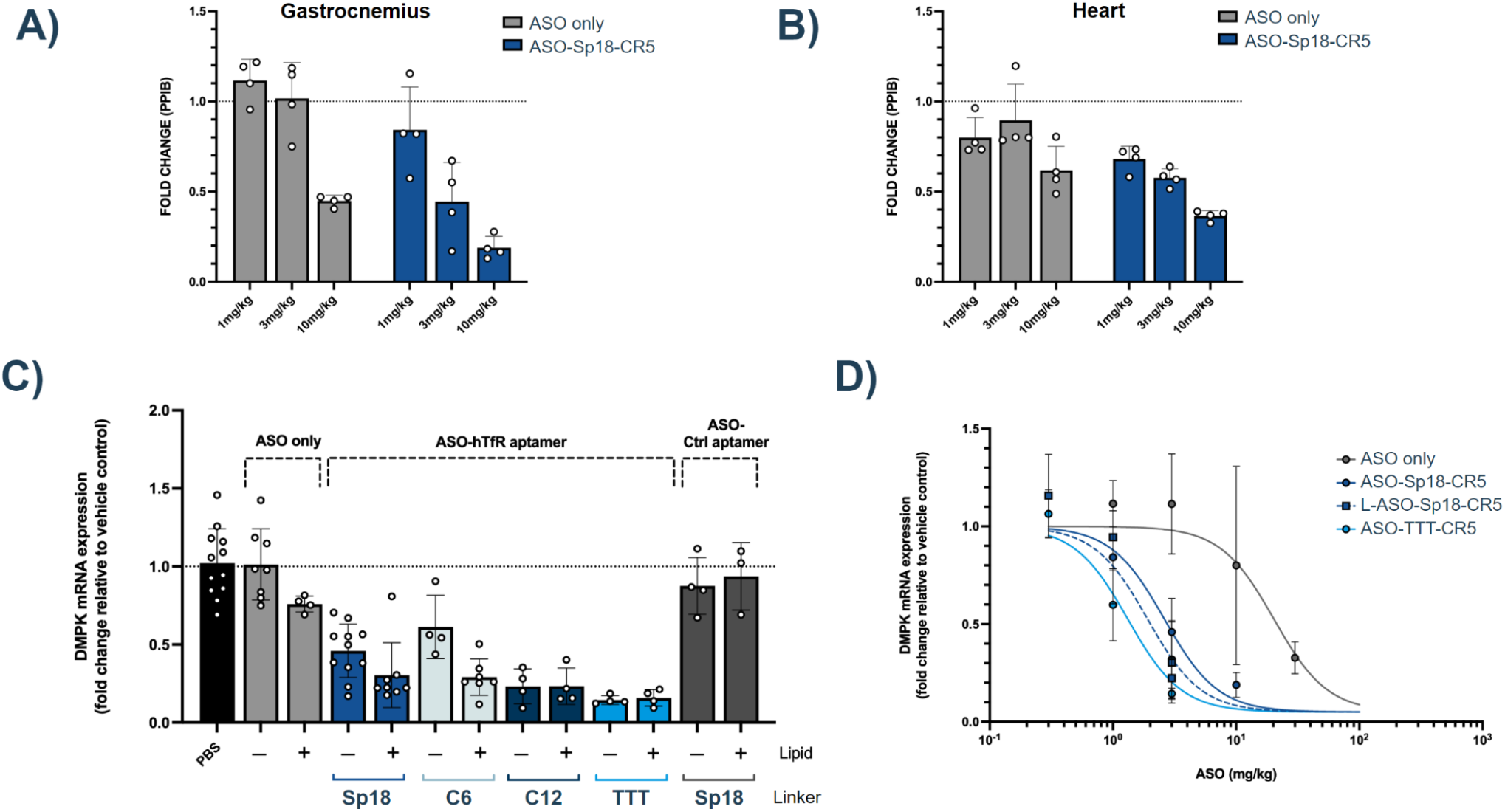
TfR aptamer mediated enhancement of ASO activity. Aptamer-ASO conjugate ASO-SP18-CR5 or ASO only was administered iv to B-hTFR1 mice on days 1, 5, and 9. Knockdown of the ASO target, DMPK, was assessed 7 days after the last dose by qPCR. PPIB served as the reference gene. **(A)** Gastrocnemius (calf) muscle **(B)** Heart muscle. **(C)** Evaluation of the effects of linker chemistry and 5’ palmitate on ASO activity in gastrocnemius muscle following iv injection of ASO only or Aptamer-ASO conjugates at 3 mg/kg and **(D)** dose-dependent knockdown of DMPK mRNA in gastrocnemius muscle following iv injection of ASO only at 3, 10, and 30 mg/kg, or Sp18 or TTT linked Aptamer-ASO conjugates at 0.3, 1, and 3 mg/kg (mass based on ASO). Animals were treated as in **(A), (B)** and **(C)**.

Follow-up experiments focused on investigating the role of the linker moiety attaching the ASO to the aptamer as well as the potential to slow clearance and enhance delivery via lipidation (17, 18). Linkers separating the aptamer and the ASO portion including, Sp18, 1,12-dodecanediol (C12), 1,6-hexanediol (C6). An additional compound was generated using three thymidine residues (TTT) as the linker, an approach which has been shown to enhance the delivery of siRNA (19). Each compound was synthesized as a single molecule in a single synthesis run using commercially available linker reagents with or without a 5’ palmitate. After deprotection and purification, a competition binding assay was used to evaluate binding activity. All compounds performed as well or better than the parent aptamer (**Supplemental Fig. 1**).

As shown in **Fig. 2C**, both linker identity and lipidation affected ASO activity. For example, when combined with an Sp18 or a C6 linker, the addition of a 5’palmitate improved DMPK knockdown activity ∼2 fold. However, for the aptamer-ASO conjugates incorporating the C12 and TTT linkers, the 5’palmitate had no additional effect, but the linker itself enhanced DMPK knockdown activity. A similar trend was observed in heart muscle (**Supplemental Fig. 2**). Importantly, in control experiments in which we lipidated just the ASO (ASO only; **Fig. 2C**), or assessed activity of a non-functional aptamer-ASO conjugate with or without lipid (ASO Ctrl aptamer; **Fig. 2C**), little to no target gene knockdown activity was observed, further supporting the role of aptamer-based TfR targeting in driving activity. Taken as a whole, these data demonstrate that aptamer targeting can enhance ASO muscle activity ∼5-6 fold and that with optimal linker chemistry or lipidation, this can be further improved to ∼9-10 fold (**Fig. 2D**)

To better understand the effect of aptamer targeting on activity, we also assessed ASO concentrations in GA tissue following treatment. Consistent with literature observations, lipidation of the ASO only had a modest, ∼1.5 fold effect on ASO tissue concentration in GA muscle (17, 18). A similar effect in tissue concentration was also observed for compounds containing the Sp18 and C6 linker which showed activity improvements with lipidation (**Fig. 3**). For compounds incorporating the C12 and TTT linkers, which did not show activity improvements following lipidation, ASO tissue concentrations were similarly not affected. Interestingly, for all compounds tested, aptamer targeting appears to have little to no effect on ASO tissue concentrations; thus, the significant improvements in activity observed for aptamer targeted compounds when compared to ASO only are not due to significant changes in ASO tissue concentrations.

**Figure 3:**
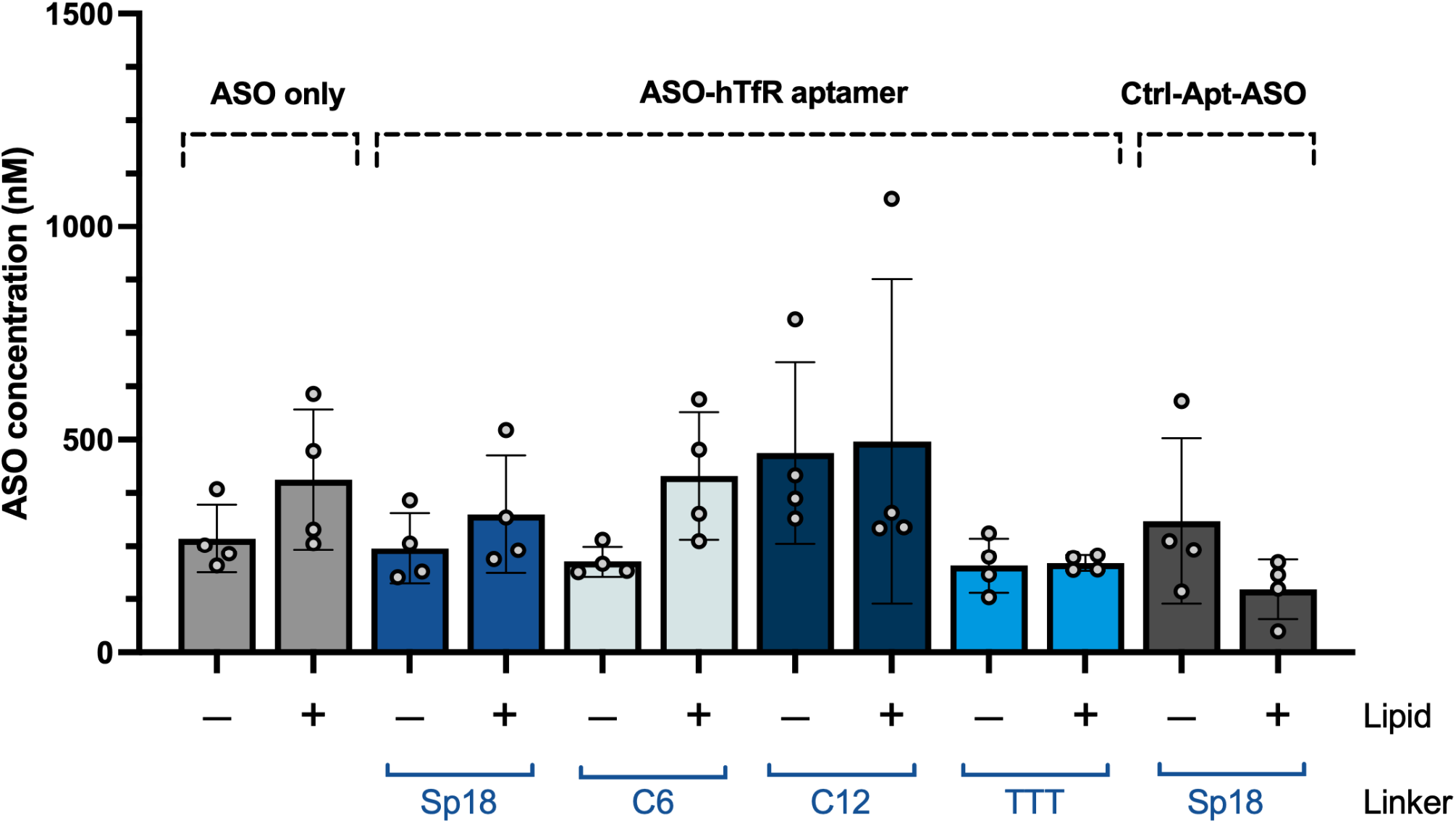
ASO concentration in gastrocnemius muscle tissue. Tissue concentrations were determined by HybELISA. Animals evaluated are the same as those shown in **Fig. 2C**.

Aptamer-ASO conjugates containing a C12 linker (ASO-C12-CR5) with or without a 5’ palmitate were further evaluated in hTfR1 knock-in mice. For these studies, mice were dosed with 3 mg/kg compound either intravenously (iv) or subcutaneously (sc) on days 1, 5, and 9, and tissues analyzed 7, 14, and 21 days after the last dose (**Fig. 4**). DMPK expression was subsequently assessed in GA, quadriceps, diaphragm, heart and liver.

**Figure 4:**
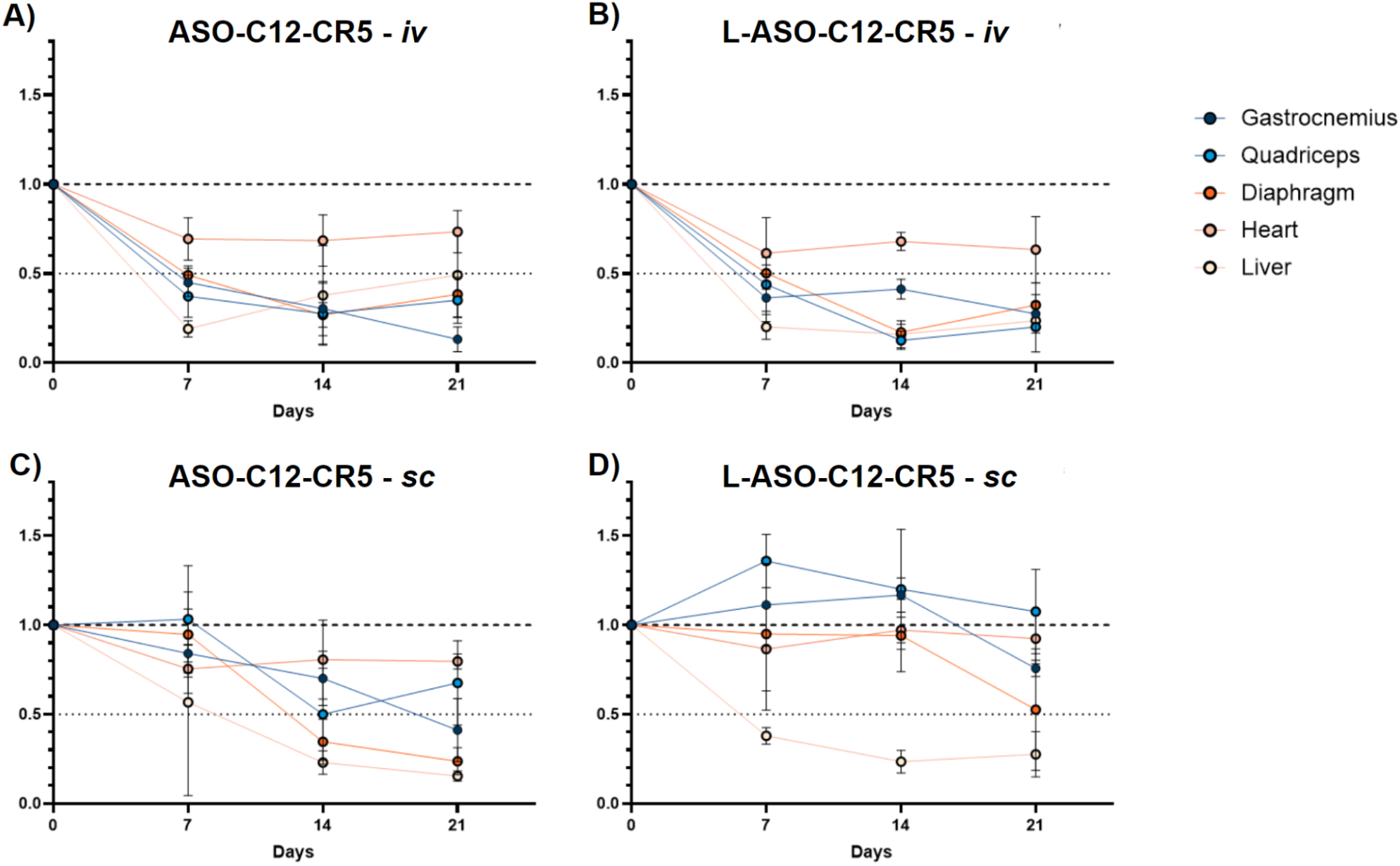
DMPK knockdown in target tissues following multidose iv or sc administration. Aptamer-ASO conjugates constructed with a C12 linker with or without a 5’ palmitate were administered at 3 mg/kg iv or sc (as indicated) on days 1, 5, and 9. Animals were euthanized 7, 14, or 21 days post dose, and the level of DMPK mRNA was determined by qPCR. PPIB served as the reference gene.

For animals treated by iv administration, target knockdown was observed at the earliest time point (7 days post dose) and continued out to the latest time point, 21 days post last dose in all tissues tested (**Fig. 4A** and **B**). In some instances, knockdown activity appears to continue to increase over the sampling period (e.g., GA muscle). As observed previously (**Fig. 2**) lipidation had little effect on compound activity with the exception of the liver where DMPK knockdown activity of the non-lipidated compound lessened over time, starting at ∼75% knockdown at day 7 post dose but rising to ∼ 50% by day 21.

For animals treated by sc administration, less activity was observed in all treatment groups at the earliest timepoints with a progressive increase in target gene knockdown over time. Interestingly, the non-lipidated compound performed better than one bearing palmitate. At 21 days post last dose, a knockdown efficiency of ∼60% was observed in GA and almost 80% in diaphragm rivaling activity observed by compounds administered iv. Similar results were observed when experiments were performed using compounds incorporating the TTT linker (ASO-TTT-CR5) and administered both iv and sc (**Supplemental Fig. 3**).

As a follow-up, we performed a single dose study in which hTfR mice were dosed with ASO-C12-CR5 at 3, 10, and 30 mg/kg, and animals were subsequently euthanized on days 7, 21, and 42. As shown in **Fig. 5**, even at the 3 mg/kg dose, ∼50% knockdown was observed in GA, quad, and diaphragm at 42 days post last dose indicating the long lasting activity of the delivered compound in muscle. However, as observed in the multidose study (**Fig. 4A**), liver activity appears to wane and returns to baseline 42 days post dose for animals treated with 3 or 10 mg/kg (**Fig. 5D**).

**Figure 5:**
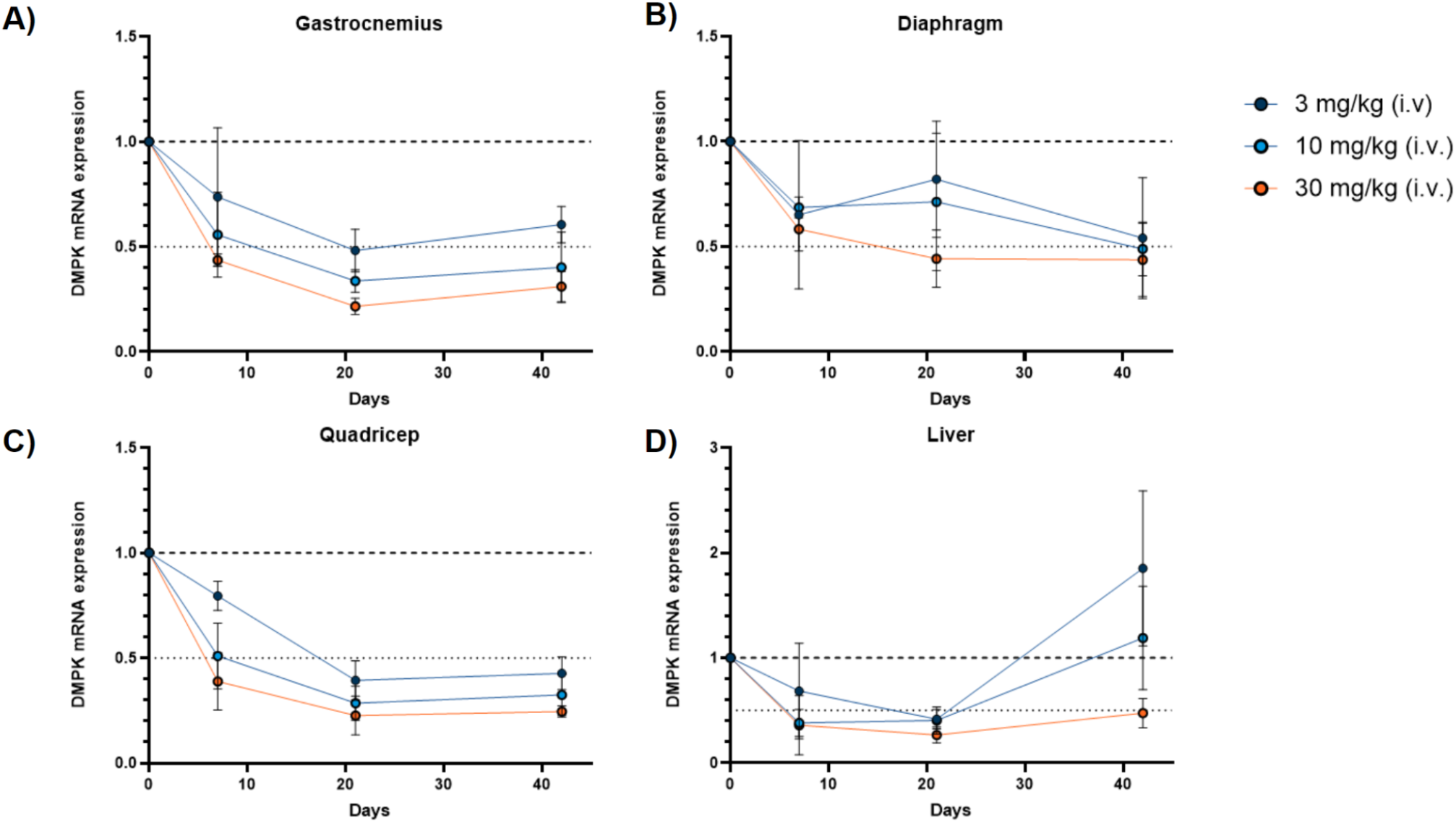
DMPK knockdown in target tissues following single iv administration. Aptamer-ASO conjugates constructed with a C12 linker without a 5’ palmitate (ASO-C12-CR5) were administered iv at 3, 10 or 30 mg/kg (mass based on ASO). Animals were euthanized 7, 21 or 42 days post dose, and the level of DMPK mRNA was determined by qPCR. PPIB served as the reference gene.

## Discussion

While antibody conjugates have led the way for extrahepatic delivery and are now being tested in the clinic (2), these protein-based compounds pose some significant limitations due to their molecular size and biological origin. Their larger size, in particular, has already led investigators to explore other options including antibody fragments (20, 21) and non-natural protein scaffolds (e.g., centyrins) (22). However, as proteins, challenges stemming from the complexity of synthesis and potential immunogenicity still remain.

The idea of using aptamers for target delivery is not new. However, many of the compounds previously described as delivery vehicles fail to function robustly (9). Moreover, many/most published aptamers are not stable enough for drug development and still susceptible to nucleases (23). Even the often employed “nuclease stable” 2’F RNA is still 50% natural RNA (the A and G residues are 2’OH). These positions are liabilities with regard to stability and potential immunogenicity and add an additional level of complexity to chemical synthesis requiring an orthogonal deprotection step that typically involves hydrogen fluoride.

Here, we have used a novel fully backbone modified aptamer which targets the human transferrin receptor and does not compete with the endogenous ligand, Tf, for binding to the receptor to potentiate making drug-like molecules. As they are fully backbone modified, such compounds are exceptionally stable (t_1/2_>100hr; (23)), non-immunogenic, and non-toxic; they are composed of the same nucleotide composition employed in extant siRNA drugs (24). When conjugated to an ASO, the resultant compound is 56 nucleotides in length, maintains binding activity, and is easily chemically synthesized, deprotected, and purified; ASO conjugates can be made in a single solid-phase synthesis run. Indeed, in pilot synthesis runs we have already made similar 56mer ASO-aptamer conjugates with ∼50% yield and ∼90% purity using “off-the-shelf” conditions (data not shown). Yields and purity will only improve with optimization.

Consistent with observations using other TfR targeting ligands, our TfR-aptamer targeted approach enhances ASO muscle activity when compared ASO alone or experiments using a non-functional aptamer control sequence (**Fig. 2C**).

Because ASO release is likely to play an important role in activity, we performed experiments focused on investigating the role of the linker moiety attaching the ASO to the aptamer. Additionally, recognizing that lipidation delays the systemic clearance of ASOs by enhancing plasma protein binding (18, 25) and can improve the activity of ASOs in muscle (17, 25), we also assessed the effects of lipidation on ASO-mediated knockdown by adding a palmitate moiety to the 5’ end of each compound. For linker chemistry, we focused on using commercially available phosphoramidite reagents which could be seamlessly integrated into a standard oligonucleotide synthesis run. This included hydrophobic linkers like C6 and C12, hydrophilic linkers like Sp18 and a DNA based linker, composed of 3 thymidine residues (TTT). Interestingly, some linkers (C12 and TTT) performed better than others (Sp18 and C6). Alternatively, some linkers (Sp18 and C6) improved by the addition of a 5’ palmitate while others (C12 and TTT) did not.

It is also interesting to note that in all cases, the increase in ASO activity in muscle tissue was not accompanied by a significant increase in ASO tissue concentration. This observation stands in contrast to the targeted delivery of siRNA in which TfR targeting results in an increase in muscle tissue siRNA concentrations (16).

The absence of a targeting effect on ASO tissue concentrations is perhaps not surprising. As a class, ASOs demonstrate broad tissue distribution as a consequence of their high PS content which does not appear to be grossly affected by the presence of the aptamer (both functional and non-function aptamer ASO conjugates show similar levels of tissue concentration to both each other and to ASO only; **Fig. 3**). In published work from Denali using a modified Fc domain to target the transferrin receptor and deliver ASOs, targeted uptake in the heart was less than 2-fold greater than that of a non-targeted ASO (see Figure 3d in (3), and drug delivered to quadriceps muscle was reduced with targeting when compared to non-targeted ASO (see Figure S3 in). Thus, for ASOs, it seems that perhaps high ASO concentrations in some tissues from non-targeted delivery may mask the effects achieved through a targeted approach. The improvements in activity that we observe with targeting ASOs therefore seems to be a consequence of cellular delivery, not increased bulk tissue concentration. That is, targeting improves cell-specific uptake compared to ASO alone leading to improved target gene knockdown in the targeted cells.

In summary, we have demonstrated that aptamers targeting the transferrin receptor can enhance ASO activity in the skeletal and cardiac muscles of human TfR1-expressing mice. This targeted delivery approach significantly streamlines the development of targeted oligonucleotide therapeutics by reducing compound synthesis to run on an oligo synthesizer using very standard and scalable conditions. Consequently, this platform offers a simpler, faster, and potentially safer alternative to protein-based delivery systems.

## Methods

### Aptamer and Aptamer-ASO Conjugate Synthesis

All aptamers and aptamer conjugates were synthesized by Synoligo Biotechnologies (Morrisville, NC) using standard solid phase nucleic acid synthesis chemistry. Following cleavage and deprotection, all compounds were HPLC purified and converted to the sodium salt before use. A full list of aptamer sequences can be found in **Supplemental Table 1**.

### Aptamer Labeling

All aptamer conjugates were generated from 5’amine modified aptamers using BP Fluor 647 TFP ester. In a typical reaction, 10 nmoles aptamer were desalted into 0.1M NaHCO3, pH 8.3 and combined with a 10-fold molar excess of dye in a final volume 100 µL. Reactions were allowed to proceed overnight at room temperature after which free dye was removed by desalting through a BioSpin 6 column (BioRad) pre-equilibrated with PBS. Dye conjugation efficiency was determined based on the ratio of 260 to 650 and the corresponding extinction coefficients. Conjugation efficiency typically proceeded to >80%.

### Flow Cytometry Binding Analysis -

Aptamer cell binding and internalization was performed on Jurkat cells in full media (RPMI with 10% FBS). Prior to use, labeled aptamers were thermally equilibrated in DPBS by heating to 80C for 5 minutes and then allowed to cool on the bench for 15 minutes. Aptamers were incubated with ∼10^5^ Jurkat cells for 1 hr in complete RPMI media supplemented with 1 µg/µl ssDNA with or without 12.5 µM hTf. Following incubation for 60 minutes at 37°C, the cells were washed three times with FACS buffer [HBSS (Hanks Buffered Saline Solution) + 1% BSA + 0.1% Na-Azide] and analyzed by flow cytometry. Dead cells were excluded by the addition of Hoechst 33342.

### Kd Determination by Bio-Layer Interferometry

Binding affinities were assessed using bio-layer interferometry (Gator® Prime,Gator Bio, Palo Alto, CA). In short, biotinylated avi-hTfR protein (Acro Biologicals) was immobilized on streptavidin probes, and probes were subsequently incubated with thermally equilibrated unlabelled aptamers in SB1T (40 mM HEPES, pH 7.5, 125 mM NaCl, 5 mM KCl, 1 mM MgCl2, 1 mM CaCl2, 0.05% Tween-20) buffer. Binding constants were determined from a global fit of the data using the equation K_D_ = k_on_ /k_off_

### In vivo Studies

All animal studies were performed in 10 – 11 week-old male B-hTFR1 mice (Biocytogen, China) and maintained in a specific pathogen free animal facility. All animal experiments were approved by the institutional animal ethics committee and the Government of India’s Committee for Control and Supervision of Experiments on Animals. Experimental animals were housed in individually ventilated mouse cages under a 12:12 hour light-dark cycle in a temperature (22 ± 3°C) and humidity-controlled (50 ± 20 %) environment with 15–20 fresh air changes per hour. Animals were provided ad libitum access to standard laboratory chow and filtered, reverse osmosis-purified water. Mice were provided appropriate structural (mouse igloo) and behavioral enrichment. Animal health and clinical signs were monitored routinely during the study period.

### Single Dose Study

Animals were dosed iv on day 1 at 3, 10, or 30 mg/kg and euthanized on day 7, 21, and 42. RNA was extracted from tissues using standard procedures and the expression level of DMPK RNA was determined by qPCR, with PPIB serving as reference.

### Multi-dose Study

Animals were dosed at 3 mg/kg (based on the mass of ASO) on days 1, 5, and 9 iv or sc as indicated. Animal groups were euthanized on day 16, 23, and 30 corresponding to 7, 14 and 21 days post last dose. RNA was extracted from tissues using standard procedures, and the expression level of DMPK RNA was determined by qPCR with PPIB serving as reference.

### RNA Extraction and Purification

Tissues were collected and placed in RNAlater (ThermoFisher) overnight at 4°C. After complete removal of RNAlater, tissues were snap frozen. For RNA extraction, frozen samples were disrupted using Trizol (Invitrogen) and a TissueLyzer III (Qiagen). RNA was recovered using Zymo Research Direct-zol-96 RNA isolation kits as per the manufacturer’s instructions.

### qPCR Analysis

First strand cDNA was prepared using iScript Reverse Transcription kits from BioRad, and DMPK and PPIB mRNA expression was then quantitated using the TaqMan gene expression assays listed in **Supplemental Table 2** with the QuantStudio 5 Real-Time PCR system (ThermoFisher). DMPK mRNA expression was normalized to PPIB and results displayed relative to ΔΔCq values calculated for PBS control animals.

### HybELISA Protocol

Quantification of ASO was performed by Hyb-ELISA as previously described (26). Probes for the detection of the ASO and the ASO-Aptamer conjugate are listed in **Supplemental Table 3**. For tissue, quantities are reported as ng/g tissue.

## Acknowledgements

Special thanks to Jeyaprakashnarayanan “JP” Seenisamy, head of Creyon Indian Operations for all that he does and to Jason Ferrone for careful reading and editing of this manuscript.

## Notes

### Competing Interest Statement

All authors are employees of Creyon Bio, Inc.

