## Supplemental Materials for "An aptamer targeting TfR1 enhances ASO delivery to muscle tissue"

### Supplement Methods

*Competition Binding Assay (flow cytometry)* - ASO-hTfR aptamer conjugate activity was evaluated by flow cytometry. Assays were performed using Jurkat cells in full media (RPMI with 10% FBS) supplemented with 12.5 µM hTf and ssDNA (1 µg/µL). Prior to initiating the competition assay, aptamers were thermally equilibrated in DPBS by heating to 80°C for 5 minutes and allowed to cool on the bench for 15 minutes. In each condition, a fixed concentration of labeled CR5 aptamer (200 nM) was mixed with unlabeled ASO-aptamer conjugate (2 µM), unlabeled parent aptamer (2 µM) to establish a positive competition control, or DPBS to generate the non-competitor (100% binding) control, before the addition of ~10^5 Jurkat cells and incubated for 60 minutes at 37°C. Jurkat cells were then washed three times with FACS buffer [HBSS (Hanks Buffered Saline Solution) + 1% BSA + 0.1% Na-azie] before being analyzed by flow cytometry. Dead cells were excluded by the addition of Hoechst 33342. Median fluorescence intensity (MFI) values for all conditions were then background subtracted to remove autofluorescence from the live cell only negative control (0% binding) and then normalized to the non-competitor labeled CR5 aptamer (100% binding) control to compare the relative binding of the ASO-aptamer conjugates versus the parental aptamer, CR5. MFI values for ASO-aptamer conjugates less than the parental unlabeled aptamer (CR5) are indicative of enhanced binding to hTfR by the ASO-hTfR aptamer conjugate, consistent with the BLI data observed for ASO-Sp18-CR5 (**Fig. 1C**).

**Supplemental Table 1: Aptamer and Aptamer Conjugate Sequences**

| Name | sequence |
|---|---|
| CR5 | (C6NH2)mCfGfGmAmCmUmUmAmCfGmUmAfGmAmAmAfGfGmUmAmAmCfGmAmUmCmAmUmUfGmUfGmUmCmCfG |
| c2.min | (C6NH2)rGrGrGrGrGrAfUfCrArAfUfCfCrArArGrGrGrAfCfCfCrGrGrArArAfCrGfCfUfCfCfCfUfUrAfCrAfCfCfCfC |
| TR14 ST1-3 | (C6NH2)fUfUfUrAfUfUfCrAfCrAfUfUfUfUfUrGrArAfUfUrGrA |
| E3 | (C6NH2)rGrGfCfUfUfUfCrGrGrGfCfUfUfUfCrGrGfCrArAfCrAfUfCrArGfCfCfCfCfUfCrArGfCfC |
| Waz | (C6NH2)rGrGrGfUfUfCfUrAfCrGrAfUrArArAfCrGrGfUfUrArAfUrGrAfUfCrArGfCfUfUrAfUrGrGfCfUrGrGfCrArGfUfUfCfCfC |
| ASO(DMPK) | +A*+5mdC*+dA*dA*T*dA*dA*dA*T*dA*5mdC*5mdC*dG*+A*+G*+G |
| L-ASO(DMPK) | +A*+5mdC*+dA*dA*T*dA*dA*dA*T*dA*5mdC*5mdC*dG*+A*+G*+G |
| ASO-Sp18-CR5 | +A*+5mdC*+dA*dA*T*dA*dA*dA*T*dA*5mdC*5mdC*dG*+A*+G*+G(Sp18)mCmGmCfGfGmAmCmUmUmAmCfGmUmAfGmAmAmAfGfGmUmAmAmCfGmAmUmCmAmUmUfGmUfGmUmCmCfGmCmG |
| ASO-C6-CR5 | +A*+5mdC*+dA*dA*T*dA*dA*dA*T*dA*5mdC*5mdC*dG*+A*+G*+G(C6)mCmGmCfGfGmAmCmUmUmAmCfGmUmAfGmAmAmAfGfGmUmAmAmCfGmAmUmCmAmUmUfGmUfGmUmCmCfGmCmG |
| ASO-C12-CR5 | +A*+5mdC*+dA*dA*T*dA*dA*dA*T*dA*5mdC*5mdC*dG*+A*+G*+G(C12)mCmGmCfGfGmAmCmUmUmAmCfGmUmAfGmAmAmAfGfGmUmAmAmCfGmAmUmCmAmUmUfGmUfGmUmCmCfGmCmG |
| ASO-TTT-CR5 | +A*+5mdC*+dA*dA*T*dA*dA*dA*T*dA*5mdC*5mdC*dG*+A*+G*+G(TTT)mCmGmCfGfGmAmCmUmUmAmCfGmUmAfGmAmAmAfGfGmUmAmAmCfGmAmUmCmAmUmUfGmUfGmUmCmCfGmCmG |
| ASO-Sp18-Ctrl | +A*+5mdC*+dA*dA*T*dA*dA*dA*T*dA*5mdC*5mdC*dG*+A*+G*+G(Sp18)fGfGmCfGmUmAfGmUfGfAmUmUmAmUfGmAmAmUmCfGmUfGmUfGmCmUmAmAmUmAmCmAmCmfGmCmC |
| L-ASO-Sp18-CR5 | (Pal)+A*+5mdC*+dA*dA*T*dA*dA*dA*T*dA*5mdC*5mdC*dG*+A*+G*+G(Sp18)mCmGmCfGfGmAmCmUmUmAmCfGmUmAfGmAmAmAfGfGmUmAmAmCfGmAmUmCmAmUmUfGmUfGmUmCmCfGmCmG |
| L-ASO-C6-CR5 | +A*+5mdC*+dA*dA*T*dA*dA*dA*T*dA*5mdC*5mdC*dG*+A*+G*+G(C6)mCmGmCfGfGmAmCmUmUmAmCfGmUmAfGmAmAmAfGfGmUmAmAmCfGmAmUmCmAmUmUfGmUfGmUmCmCfGmCmG |
| L-ASO-C12-CR5 | (Pal)+A*+5mdC*+dA*dA*T*dA*dA*dA*T*dA*5mdC*5mdC*dG*+A*+G*+G(C12)mCmGmCfGfGmAmCmUmUmAmCfGmUmAfGmAmAmAfGfGmUmAmAmCfGmAmUmCmAmUmUfGmUfGmUmCmCfGmCmG |
| L-ASO-TTT-CR5 | (Pal)+A*+5mdC*+dA*dA*T*dA*dA*dA*T*dA*5mdC*5mdC*dG*+A*+G*+G(TTT)mCmGmCfGfGmAmCmUmUmAmCfGmUmAfGmAmAmAfGfGmUmAmAmCfGmAmUmCmAmUmUfGmUfGmUmCmCfGmCmG |
| L-ASO-Sp18-Ctrl | (Pal)+A*+5mdC*+dA*dA*T*dA*dA*dA*T*dA*5mdC*5mdC*dG*+A*+G*+G(Sp18)fGfGmCfGmUmAfGmUfGfAmUmUmAmUfGmAmAmUmCfGmUfGmUfGmCmUmAmAmUmAmCmAmCmfGmCmC |

Where “*” = phosphorothioate, “+” = LNA, “m” = 2’Omethyl, “f” = 2’F, “d” = 2’deoxy, “r” 2’OH, (Sp18) = hexadecyl ethyleneglycol, (C12) = 1,12-dodecanediol, (C6) = 1,6-hexanediol (C6) and (TTT) = three thymidine residues.

**Supplemental Table 2: TaqMan qPCR probes**

| Gene target | TaqMan assay ID# (ThermoFisher) |
|---|---|
| DMPK | Mm00446261_m1 |
| PPIB | Mm00478295_m1 |

**Supplemental Table 3: HybELISA  probes**

| **Probe ID** | **Probe Sequences** |
|---|---|
| CR-PP-01811 | B-+5MeC+5MeCT+5MeC+GG+T+A |
| CR-PP-01812 | +T+TT+A+TT+G+T-Am6-DIG |

where B = biotin, +N = LNA, Am6 = C6=hexylamine, Dig = dioxygenin

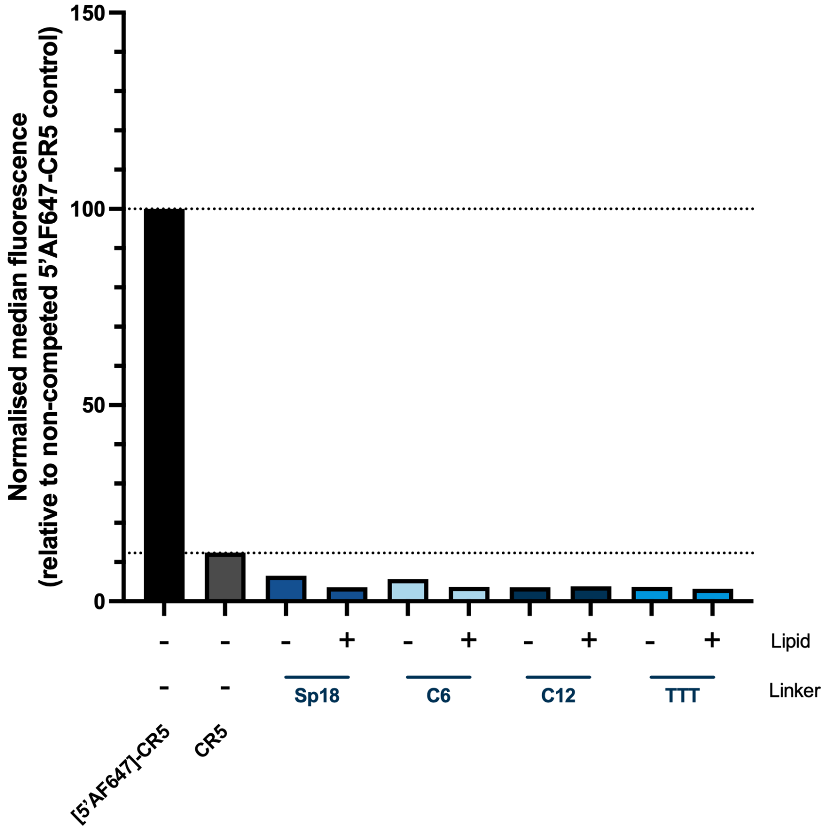


**Supplemental Figure 1.** Competition binding of ASO-hTfR aptamer conjugates versus CR5 hTfR aptamer.

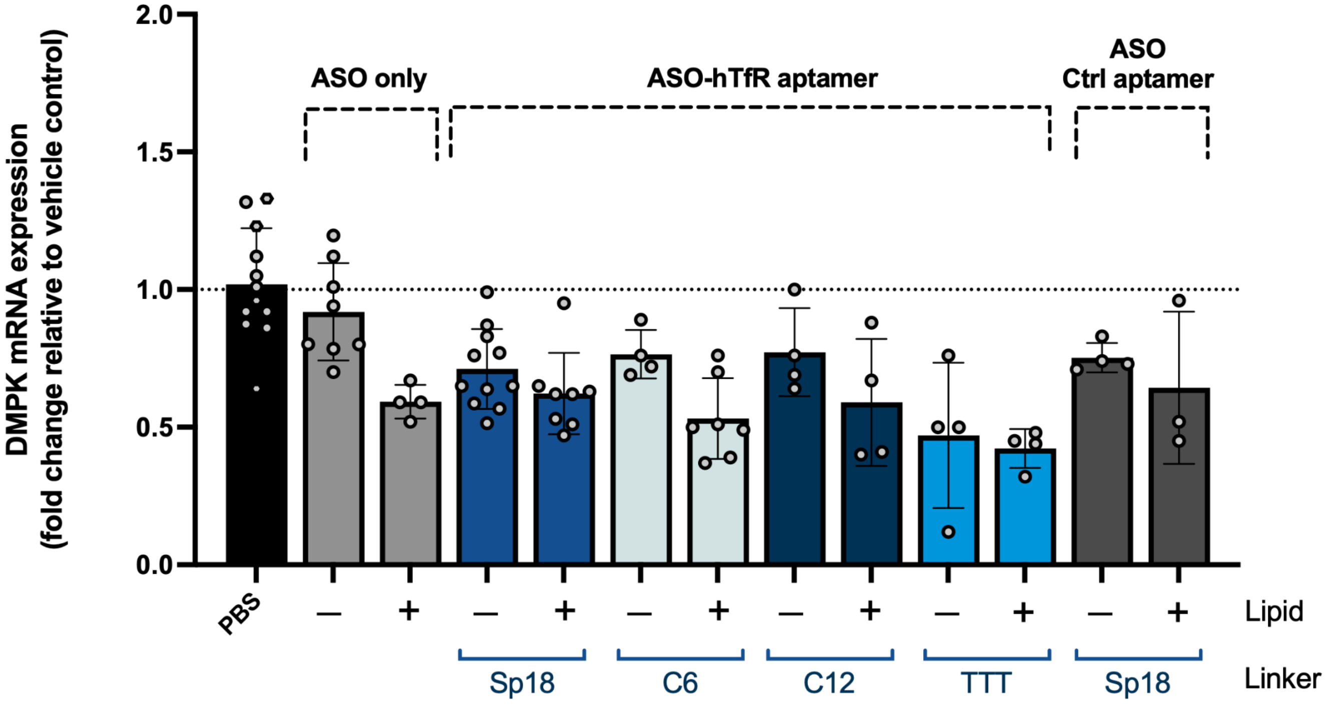


**Supplemental Figure 2. Effects of linker chemistry and 5' palmitate on ASO activity in cardiac tissue.** ASO-aptamer conjugates or ASO only was administered iv to B-hTFR1 mice on days 1, 5, and 9. Knockdown of the ASO target, DMPK, was assessed 7 days after the last dose by qPCR. PPIB served as the reference gene.

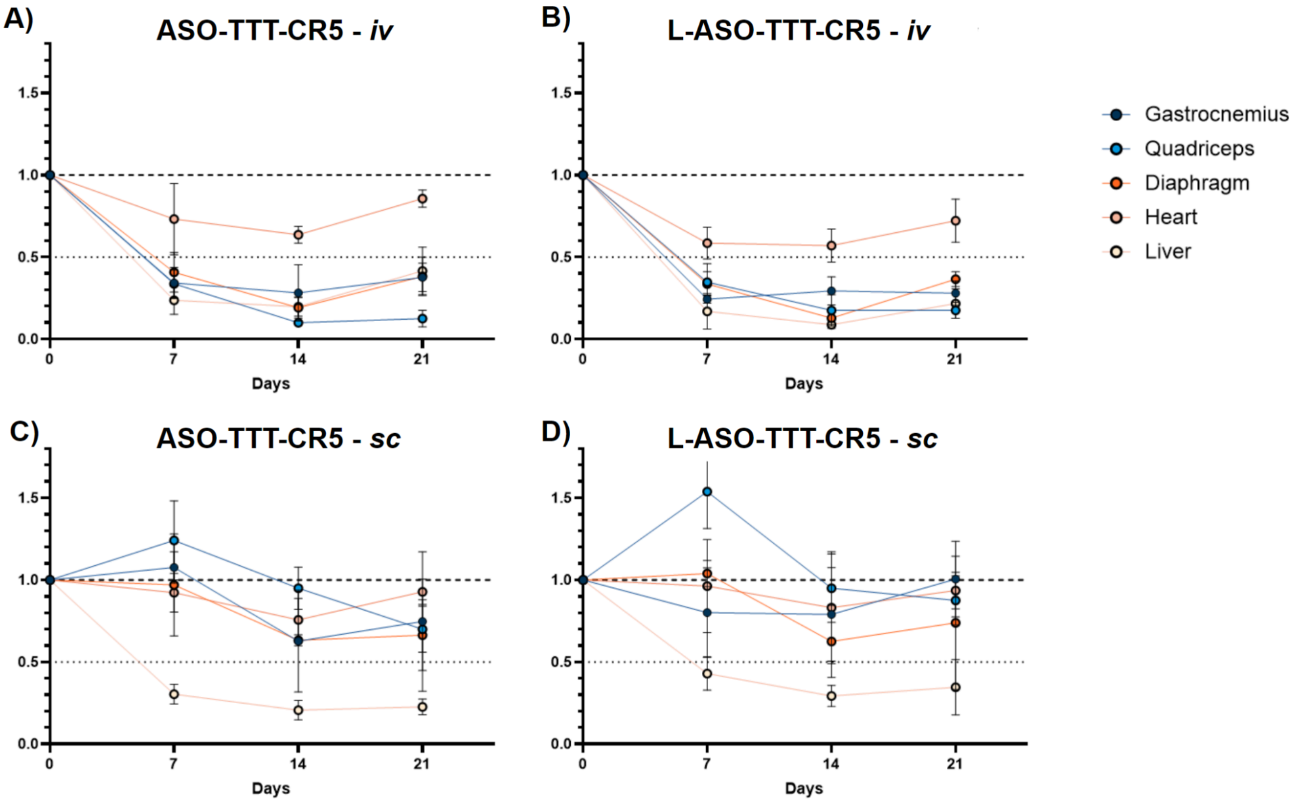


**Supplemental Figure 3. DMPK knockdown in target tissues following single iv administration.** Aptamer-ASO conjugates constructed with a TTT linker with and without a 5' palmitate (ASO-TTT-CR5 and L-ASO-TTT-CR5) were administered iv at 3 mg/kg (mass based on ASO) on days 1, 5, and 9. Animals were euthanized 7, 14 or 21 days after the last dose, and the level of DMPK mRNA was determined by qPCR. PPIB served as the reference gene.
